# Assessment of Antimicrobial Therapy in Eradicating *Chlamydia muridarum* in Research Mice: Immune Status and its Impact on Outcomes

**DOI:** 10.1101/2024.06.28.600682

**Authors:** Michael B Palillo, Sebastian E Carrasco, Noah Mishkin, Jack A Palillo, Denise B Lynch, Samira Lawton, Mert Aydin, Anthony Mourino, Neil S Lipman, Rodolfo J Ricart Arbona

**Affiliations:** Tri-Institutional Training Program in Laboratory Animal Medicine and Science, Memorial Sloan Kettering Cancer Center, Weill Cornell Medicine, and The Rockefeller University, New York, NY; Center for Comparative Medicine and Pathology, Memorial Sloan Kettering Cancer Center and Weill Cornell Medicine, New York, NY; Neurological Clinical Research Institute, Massachusetts General Hospital, Boston, MA, United States; The Jackson Laboratory, Bar Harbor, Maine; One Codex, San Francisco, California; Transnetyx, Cordova, Tennessee

## Abstract

Chlamydia muridarum (Cm) is a moderately prevalent, gram-negative, intracellular bacterium that affects laboratory mice, causing subclinical to severe disease, depending on the host’s immune status. The effectiveness of various antibiotic regimens aimed at eradicating Cm in both immunodeficient and immunocompetent laboratory mice was evaluated. NSG mice were cohoused with Cm-shedding BALB/cJ mice for 14 days to simulate natural exposure. Four groups of 8 infected NSG mice were treated for 7 days with either 0.08% sulfamethoxazole and 0.016% trimethoprim (TMS) in water, 0.0625% doxycycline in feed, 0.124%/0.025% TMS in feed, or 0.12% amoxicillin in feed. A control group was provided standard water and feed. The impact of treatment on gastrointestinal microbiota (GM) was performed using next-generation shotgun sequencing on the last day of treatment. TMS and Amoxicillin had negligible effects on GM, while doxycycline had the largest effect. All antibiotic treated NSG mice exhibited clinical disease, including dehydration, hunched posture, >20% weight loss, and dyspnea, leading to euthanasia 21-40 days post-treatment (32.6 ± 4.2 days; mean ± SD). Untreated controls were euthanized 14-33 days post-exposure (23.75 ± 5.9 days). All mice were fecal PCR positive for Cm at euthanasia. Histological evaluation revealed multifocal histiocytic and neutrophilic bronchointerstitial pneumonia and/or bronchiolitis featuring prominent intralesional chlamydial inclusion bodies in all mice. Subsequently, groups of 8 C57BL/6J, BALB/cJ, NOD.SCID, and NSG mice infected with Cm were treated with 0.124%/0.025% TMS in feed for 7 (BALB/cJ and C57BL/6J) or 21 days (NSG and NOD.SCID). All immunocompetent and NOD.SCID mice were negative for Cm by PCR 14 days post-treatment, remained clinically normal and had no evidence of Cm infection at necropsy, all NSG mice remained Cm positive and were euthanized. While these findings highlight the difficulties in eradicating Cm from highly immunodeficient mice, eradication of Cm from immunocompetent or moderately immunocompromised mice with antibiotics is feasible.

## Introduction

*Chlamydia muridarum* (Cm) is a gram-negative, obligate, intracellular bacterial pathogen of mice.^8,44^ Cm was initially misidentified as a virus in the 1930s when it was first isolated from research mice exhibiting respiratory illness.^17,24,52^ Subsequently, it was shown to be a bacterium and was reclassified as a *Chlamydia trachomatis* biovar and most recently as Cm following molecular analysis.^19,54^ Cm is the only natural chlamydial pathogen of mice and due to its similarity to *C. trachomatis* it has been used as the principal murine model of human *C. trachomatis* infection.^52^

Cm was recently isolated from and shown to be prevalent and globally distributed among research mouse colonies.^44^ Of 58 groups of mice imported from 39 academic institutions, 33% tested positive for Cm upon arrival, and 16% of a 900-diagnostic sample cohort and 14% of 120 institutions submitting approximately 11,000 murine microbiota samples had evidence of Cm.^44^ Cm infection, through its presumptive fecal-oral route of transmission of its metabolically inactive extracellular infectious elementary bodies (EBs), has been shown to cause significant disease, including severe dyspnea, pneumonia and mortality in highly immunodeficient NSG mice, and less severe pulmonary and renal lesions in genetically engineered mouse (GEM) strains with select immune system alterations.^43,44,66^ Given Cm’s ability to induce a robust immune response, cause pathology in immunocompromised mice and persistent infection in both immunocompromised and immunocompetent mice, we continue to propose that screening for Cm be routinely conducted and the bacterium excluded from research mouse colonies.^43,44,66^

Treatment of naturally infected Cm-colonized research mouse colonies has not been explored. However, select antibiotics have been used successfully in experimental Cm infections and therapies are available for treatment of chlamydia in other species, both of which can aid in the development of an effective therapeutic regimen.^6,14,31,50,51,60,67,70,71^ Developing a therapeutic strategy to eradicate Cm presents a formidable challenge, particularly in ensuring 100% effectiveness when treating an entire research mouse colony, as a treatment failure in even a small number of animals poses a risk of reinfection of animals which have been successfully treated. Adding to the complexity is the reality that many GEM models may lack some or all of the immune system components necessary to aid in Cm clearance.

Doxycycline administered at 10 to 100 mg/kg for 7 to 45 days is the treatment of choice for chlamydial infections of various animal species, including mice experimentally infected with Cm.^6,14,60,67,68,70^ While doxycycline has shown to be effective in laboratory mice, the high prevalence of Tet-on/Tet-Off systems for inducible gene expression makes it use undesirable in some research mouse colonies.^15,62^ Azithromycin is the first-line therapeutic used in humans, given that effective chlamydial eradication requires only a single oral 10 mg/kg dose.^71^ However in murine models inoculated with Cm, multiday azithromycin treatment regimens employing dosages up to 8 times higher than that used in humans yielded inconsistent results.^6,51^ Following experimental intravaginal Cm infection, amoxicillin (2 to 40 mg/kg for 7 days) was shown to be ineffective, although it is used to treat *C. trachomatis*-infected pregnant woman.^50,71^ However, amoxicillin, which is available as a compounded mouse feed at concentrations that yield higher doses than those used to treat experimental infections, may be a useful therapeutic option. Trimethoprim sulfamethoxazole (TMS) has been used to treat chlamydial infections in humans and would be desirable if effective against Cm in mice because it is commercially available as a compounded feed.^11,29,36^

Consideration must also be given to the impact of antibiotics on the microbiome when administered in the research setting. The gastrointestinal microbiota (GM) can be easily altered by antibiotic administration, leading to shifts in microbial composition that can influence the host in various ways, confounding various research models.^21,32,34,38,59^ Doxycycline and amoxicillin have been shown to induce major permanent shifts in the murine GM.^10,32,37,64^ Conversely, TMS administered in the mouse’s drinking water for 14 days causes minimal to no alterations in GM richness and β-diversity when assayed 4 weeks after antibiotic administration.^33^

Most chlamydial treatment protocols involve treatment of individual patients; however, a modern vivarium can house tens of thousands of mice requiring treatment. Therefore, in addition to being efficacious, the ideal therapeutic would be easy to administer. Administering therapeutics in feed and water would offer significant advantages when treating Cm-infected mouse colonies.^42,55,57^ The goal of the current study was to evaluate the effectiveness of various antibiotic regimens to treat Cm-infected mice with varying levels of immunocompetence to identify a highly effective, safe, easy to administer therapeutic, while minimally impacting the host’s microbiome.

## Materials and Methods

### Experimental design

#### Pilot study

A controlled antibiotic efficacy study was conducted to determine if Cm can be eradicated from outbred immunocompetent Swiss Webster mice (SW) utilizing medicated feed or water. Treatment regimens were guided by a literature review and pilot study results would guide treatment protocols used in subsequent experiments.^6,14,23,40,50,51,67^ The SW mice had been utilized as soiled bedding sentinels, were confirmed positive for Cm by PCR, and were randomly assigned to 1 of 8 treatment groups of 8 mice, housed in 2 cages each containing 4 mice. Groups included: *Control* (untreated); *Amoxicillin (Amoxi) Diet*; *Doxycycline (Doxy) Diet*; *Doxy Water*; *Enrofloxacin (Enro) Water*; *Trimethoprim-Sulfamethoxazole (TMS) Diet*; *TMS Diet-Short*, and *TMS Water*. Detailed treatment product and dosing information are provided in Table 1. The *Control* group was provided an unmedicated, irradiated (base) diet (LabDiet 5053, PMI, St. Louis, MO) ad libitum for the entirety of the study. *TMS Diet-Short* group received medicated feed for only 3 days. All other groups received medicated feed or water for 7 days. At the end of the treatment period, mice were provided the base diet and RO acidified water. A fecal pellet was harvested from each mouse and pooled by treatment group for Cm testing by PCR at 7-, 14-, and 58-days post-treatment (DPT). A naive male NOD.Cg-Prkdc^scid^Il2rg^tm1Wjl^/SzJ (NSG) mouse was added to each cage in all treatment groups at 14 DPT to serve as a contact sentinel highly susceptible to clinical disease, if a negative post-treatment PCR result on 7 DPT was confirmed. Treated mice were euthanized if the post-treatment PCR result was positive at any timepoint or at 58 DPT, the study endpoint. A gross necropsy was performed at euthanasia.

**Table 1.** Antibiotic feed and water therapies utilized in all study arms.

| **Antibiotic** | **Estimated\***<br>**Daily Dose**<br>**(mg/kg)** | **Delivery**<br>**Method** | **Abbreviation(s)** | **Formulation** | **Company,**<br>**City, State** |
| --- | --- | --- | --- | --- | --- |
| Amoxicillin | 228.0 | Diet | Amoxi Diet | PicoLab Mouse Diet 5053 with 0.12% Amoxicillin | TestDiet, St. Louis, MO |
| Doxycycline | 118.8 | Diet | Doxy Diet | Teklad Global 18% Protein Rodent Diet with 625 ppm doxycycline | Teklad Custom Diets, Madison, WI |
| Doxycycline | 12.7 | Water | Doxy Water | 25mg (5 mL) doxycycline monohydrate in 500mL of drinking water | Lupin Pharmaceuticals Inc., Baltimore MD |
| Enrofloxacin | 102.3 | Water | Enro Water | 200 mg (2 mL) enrofloxacin in 500mL drinking water | Bayer, Shawnee, Mission, KS |
| Sulfamethoxazole (S) /Trimethoprim (T) | 235.6 (S)<br>47.5(T) | Diet | TMS Diet-Short (3-day course)<br>TMS Diet (7-day course)<br>TMS Diet-Long (21-day course) | PicoLab Mouse Diet 5053 with 0.124% Sulfamethoxazole/ 0.025% Trimethoprim | TestDiet, St. Louis, MO |
| Sulfamethoxazole (S) /Trimethoprim (T) | 201.3 (S)<br>40.3 (T) | Water | TMS Water | 400 mg sulfamethoxazole, 80 mg trimethoprim solution (10.0 mL) in 500mL of drinking water | Pharmaceutical Associates, Inc., Greenville, SC |

\* Based on a 30-gram mouse ingesting 5.7 gm of food or 7.7 mL of water daily. (Bachmanov 2002)

#### Cm-infected NSG mice treatment study

A controlled antibiotic efficacy study was then conducted to determine if Cm can be eradicated from highly susceptible NSG mice using medicated feed or water.^66^ Forty-eight Cm-negative (confirmed by PCR) female NSG mice housed 4 mice per cage were acclimated for 14 days. Twelve female Cm-shedding BALB/cJ (C) mice from a Cm positive colony (details below) were randomly selected and cohoused with each cage of 4 female NSG mice for a 14-day exposure period to facilitate natural transmission of Cm. On day 14, the Cm-infected C mice were removed from each cage and the remaining NSG mice were transferred to clean cages, confirmed Cm positive by fecal PCR pooled by cage, and randomly assigned to 1 of 6 treatment groups, each consisting of 8 mice housed 4 per cage. Study groups included: *Control* (untreated); *Amoxi Diet*; *Doxy Diet*; *TMS Diet*; *TMS Diet-Long* or *TMS Water*. The *Control* group was provided unmedicated base diet for the entirety of the study. *TMS Diet-Long* group received medicated feed for 21 days. All other groups were treated for 7 days. At the end of the treatment period mice were provided unmedicated base diet and acidified water and were retested for Cm by PCR at euthanasia. Mice were monitored throughout the study as described below and euthanized when humane criteria were reached at which time a fecal pellet was collected from each mouse and pooled by cage for Cm testing by PCR. Mice in the *TMS Diet-Long* group were retested for Cm by fecal PCR at 14 DPT and euthanized if Cm positive, even though the mice had not reached humane endpoint criteria. To assess changes in GM diversity, feces were collected from 2 NSG mice per cage from all groups, except *TMS Diet-Long*, at 3 timepoints: before Cm colonization (after acclimatization before cohousing), 14 days post-cohousing (DPC), and on the final day of treatment (7 DPT). Fecal samples were collected between 7 am and 9 am.

#### Inbred immunocompetent & NOD.SCID treatment study

Following treatment failures in the prior study, an antibiotic efficacy study was conducted to investigate whether differences in host immune status can influence the effectiveness of therapy. *TMS Diet* was selected for this study based on results from the Cm-infected NSG mice treatment study, the expectation that TMS would have the least effect of the antibiotics tested on the microbiota, and the ease of administration.^33^ Female, Cm-free (confirmed by PCR), C, C57BL/6J (B6) and NOD.Cg-*Prkdc^scid^*/J (NOD.SCID) mice (n=8/strain) were randomized, housed 4 mice per cage by strain, and were infected with Cm as described previously. NOD.SCID mice were utilized as while they are immunocompromised, however they are less immunodeficient than NSG mice.^61^ C and B6 mice were treated with *TMS Diet* for 7 days while the immunocompromised NOD.SCID mice were treated for 21 days (*TMS Diet-L*). At the end of the treatment period, mice were provided base feed. A fecal pellet was harvested from each mouse and pooled by treatment group for Cm testing by PCR at 7-, 14- and 40-DPT. A naive female NSG mouse was added to each cage in all treatment groups, following a negative post-treatment PCR result 7 DPT. Two cohoused Cm-infected NOD.SCID mice were maintained as an untreated positive control group. Mice were monitored daily and euthanized 40 DPT, or earlier, if humane endpoints were met.

### Animals

Six- to 12-month-old female SW mice (*n* = 64, Tac:SW, Taconic Biosciences, Germantown, NY) and 6 to 12-month-old male NSG (*n* = 16, NOD.Cg-*Prkdc^scid^Il2rg^tm1Wjl^*/SzJ, The Jackson Laboratory, Bar Harbor, ME) were used in the *Pilot Study*. Five- to 8-wk-old female NSG mice (*n* = 48; The Jackson Laboratory) were used in the *Cm-infected NSG treatment study*. Five to 8-wk-old female C mice (*n* = 8, The Jackson Laboratory), 5 to 8-wk-old female B6 mice (*n* = 8, The Jackson Laboratory), 3 to 5-wk-old male NSG (*n* = 8) and 5 to 8-wk-old NOD.SCID (*n* = 10, The Jackson Laboratory) were used in the *Inbred immunocompetent & NOD.SCID treatment study*. Cm shedding female C mice (*n* = 19; age 38 to 49-wk-old) were used to naturally transmit Cm to all naïve mice in the *Cm infected NSG treatment study* and the *Inbred immunocompetent & NOD.SCID treatment study.* All mice were individually ear-tagged, assigned a distinct numerical identifier, and randomly assigned to appropriate groups using an open-sourced online randomizer (https://www.random.org/lists/). All mice, except those used for the SW mice used in the *Pilot study* were free of mouse hepatitis virus (MHV), Sendai virus, mouse parvovirus (MPV), minute virus of mice (MVM), murine norovirus (MNV), murine astrovirus 2 (MuAstV2), pneumonia virus of mice (PVM), Theiler meningoencephalitis virus (TMEV), epizootic diarrhea of infant mice (mouse rotavirus, EDIM), reovirus type 3, lymphocytic choriomeningitis virus (LCMV), K virus, mouse adenovirus 1 and 2 (MAD 1/2), polyoma virus, murine cytomegalovirus (MCMV), mouse thymic virus (MTV), Hantaan virus, murine chaphamaparvovirus-1(MuCPV), *Mycoplasma pulmonis*, CAR *bacillus* (*Filobacterium rodentium*), *Chlamydia muridarum*, *Citrobacter rodentium, Rodentibacter pneumotropicus, Helicobacter* spp., segmented filamentous bacterium (SFB), *Salmonella* spp., *Streptococcus pneumoniae*, Beta-hemolytic *Streptococcus* spp., *Streptobacillus moniliformis, Clostridium piliforme, Corynebacterium bovis, Corynebacterium kutscheri, Staphylococcus aureus, Klebsiella pneumoniae, Klebsiella oxytoca*, *Pseudomonas aeruginosa, Myobia musculi*, *Myocoptes musculinis*, *Radfordia affinis*, *Syphacia* spp. *Aspiculuris* spp., *Demodex musculi*, *Pneumocystis* spp, *Giardia muris*, *Spironucleus muris*, *Entamoeba muris*, *Tritrichomonas muris*, and *Encephalitozoon cuniculi* when studies were initiated.

### Husbandry and housing

All mice were maintained in individually ventilated, autoclaved, polysufone cages with stainless-steel wire bar lids and filter tops (# 19, Thoren Caging Systems; Hazelton, PA) on autoclaved aspen-chip bedding (PWI Industries, Quebec, Canada) at a density up to 5 mice per cage. Each cage was provided with a bag constructed of Glatfelter paper containing 6 g of crinkled paper strips (EnviroPak^®^, WF Fisher and Son, Branchburg, NJ) and a 2-inch square of pulped virgin cotton fiber (Nestlet^®^, Ancare, Bellmore, NY) for enrichment. Each cage (including cage bottom, bedding, wire bar lid and water bottle) was changed weekly within a class II, type A2 biological safety cabinet (LabGard S602-500, Nuaire, Plymouth, Mn). All mice were fed a natural ingredient, closed source, autoclavable diet (5KA1, LabDiet, Richmond, VA) and provided ad libitum autoclaved reverse osmosis acidified (pH 2.5 to 2.8 with hydrochloric acid) water in polyphenylsulfone bottles with autoclaved stainless-steel caps and sipper tubes (Tecniplast, West Chester, PA), unless provided medicated feed or water. All cages were housed within a dedicated, restricted-access cubicle, which was maintained on a 12:12-h light:dark (6 AM on:6 PM off) cycle, relative humidity of 30% to 70%, and room temperature of 72 ± 2°F (22 ± 1°C).

The animal care and use program at Memorial Sloan Kettering Cancer Center (MSK) is accredited by AAALAC International and all animals are maintained in accordance with the recommendations provided in the Guide.^5^All animal use described in this investigation was approved by MSK’s IACUC in agreement with AALAS position statements on the Humane Care and Use of Laboratory Animals and Alleviating Pain and Distress in Laboratory Animals.

### Generation of Cm shedding mice

C mice were experimentally inoculated with Cm 8 to 11 months prior to the initiation of this study via oral gavage with a wild-type stock (Cm field strain isolated from clinically affected NSG mice at MSK) at a concentration of 2.72×10^3^ IFU in 100uL of sucrose-phosphate-glutamic acid buffer (SPG, pH 7.2) using a sterile orogastric gavage needle (22g x 38.1mm, Cadence Science, Cranston, RI).^44^ Mice were confirmed Cm positive by fecal PCR following initial infection and on the day of cohousing.

### Medicated feed and water

Details related to antibiotic feed and water with product information are provided in Table 1. Medicated water was prepared and provided in autoclaved polyphenylsulfone bottles with stainless-steel caps and sipper tubes, containing reverse osmosis water acidified to a pH of 2.5 to 2.8 with hydrochloric acid. While we did not measure food or water consumption, estimated ingested dosages of medicated feed and water were determined using established feed and water consumption data for a 30-gram mouse of 5.7 gm and 7.7 ml per day, respectively.^3^

### Clinical evaluation, weight assessments and humane endpoint criteria

All mice were observed cage side daily for morbidity, including but not limited to lethargy, hunched posture, piloerection, change in body condition, dehydration, dyspnea or cyanosis. All mice were weighed prior to study initiation to provide a baseline weight. If clinical signs of disease developed, mice were weighed daily and euthanized when they lost ≥20% of their initial body weight.

### Fecal collection

Fecal pellets for PCR or GM analysis were collected from each mouse. Mice were lifted by the base of the tail and allowed to grasp the wire bar lid while a sterile 1.5-mL microfuge tube (PCR) or collection kit tube (GM analysis) was held below the anus to allow feces to fall directly into the tube. If the mouse did not defecate within a 30 sec period, it was returned to the cage and allowed to rest for at least 2 min before collection was reattempted until a sample was produced. A single pellet was collected for PCR analysis and 2 fecal pellets per animal were collected for GM analysis. PCR samples were pooled by cage and stored at −80 °C (−112 °F) until analyzed.

### Pathology

Mice were euthanized by carbon dioxide (CO_2_) asphyxiation in accordance with recommendations.^1^ Euthanized mice were submitted for a complete necropsy. Macroscopic changes in tissues were recorded and tissues were collected and fixed in 10% neutral buffered formalin for at least 72 hours. After fixation, the skull was decalcified in a formic acid and formaldehyde solution (Surgipath Decalcifier I, Leica Biosystems, Deer Park, IL). Tissues were then processed in ethanol and xylene, embedded in paraffin (Leica ASP6025 processor, Leica Biosystems), sectioned (5-μm-thick) and stained with hematoxylin and eosin (H&E) or processed for immunohistochemistry (IHC). In the *Cm-infected NSG mice treatment study* H&E stained lung sections were evaluated for evidence of Cm-mediated pneumonia.^66^ In the *Inbred immunocompetent & NOD.SCID treatment study* H&E stained lung, small intestine, large intestine, uterus, vagina, urinary bladder and head (including brain, eye, ears, pituitary, nasal and oral cavities, teeth, soft tissues, skin) sections from Control NOD.SCID mice were evaluated.

Formalin-fixed, paraffin-embedded tissues (lung, small intestine, large intestine, uterus, vagina, and urinary bladder) harvested from the *Inbred immunocompetent & NOD.SCID treatment study* mice were stained for Cm major outer membrane protein (MOMP) using an automated staining platform (Leica Bond RX, Leica Biosystems). After deparaffinization and heat-induced epitope retrieval in citrate buffer at pH 6.0, the primary antibody against chlamydial MOMP (NB100-65054, Novus Biologicals, Centennial, CO) was applied at a dilution of 1:500. A rabbit anti-goat secondary antibody (Cat. No. BA-5000, Vector Laboratories, Burlingame, CA) and a polymer detection system (DS9800, Novocastra Bond Polymer Refine Detection, Leica Biosystems) was then applied to the tissues. The 3,3’-diaminobenzidine tetrachloride (DAB) chromogen was used, and the sections were counterstained with hematoxylin and examined by light microscopy. Intestines and reproductive tracts from NSG or TLR3-deficient mice infected with *Chlamydia muridarum* strain Nigg were used as positive control.^12^ Positive MOMP hybridization was identified as discrete, punctate chromogenic brown dots under bright field microscopy. A board-certified veterinary pathologist (SEC) performed all histologic analyses.

### *Chlamydia muridarum* qPCR assay

DNA and RNA were copurified from feces or cage swab samples using the Qiagen DNeasy 96 blood and tissue kit (Qiagen, Hilden, Germany). Nucleic acid extraction was performed using the manufacturer’s recommended protocol, “Purification of Total DNA from Animal Tissues”, with the following buffer volume modifications: 300µL of Buffer ATL + Proteinase K, 600µL of buffer AL + EtOH, and 600µL of lysate were added to individual wells of the extraction plate. Washes were performed with 600µL of buffers AW1 and AW2. Final elution volume was 150µL of buffer AE.

A probe-based PCR assay was designed using IDT’s PrimerQuest Tool (IDT, Coralville, IA) based on the 16s rRNA sequence of *Chlamydia muridarum*, Nigg strain (Accession NR_074982.1, located in the National Center for Biotechnology Information database). Primer and Probe sequences generated from the PrimerQuest Tool were checked for specificity using NCBI’s BLAST (Basic Local Alignment Search Tool). The probe was labeled with FAM and quenched with ZEN and Iowa Black FQ (IDT, Coralville, IA). Primer names, followed by associated sequences are as follows: *C. muridarum*_For (GTGATGAAGGCTCTAGGGTTG); *C. muridarum*_Rev (GAGTTAGCCGGTGCTTCTTTA), *C. muridarum*_Probe (TACCCGTTGGATTTGAGCGTACCA).

The Cm PCR assay was validated by generating a standard curve. Positive PCR amplicons derived from known Cm positive fecal samples were combined to a volume of 25µl and purified by pipetting using Diffinity Rapid Tips (Diffinity Genomics, West Chester, PA). Five, 10-fold serial dilutions were run in triplicate producing the following values calculated by Bio-Rad CFX Maestro Software (Bio-Rad, Hercules, CA): E=97.6%, R^2=.995, Slope=-3.381, y-intercept=41.781. To estimate copy numbers, serial dilutions were quantified using a Qubit 2.0 Fluorometer and Qubit dsDNA Quantitation, High Sensitivity Assay (Thermo Fisher Scientific). The concentration values were calculated using the Thermo Fisher DNA Copy Number and Dilution Calculator with default settings for molar mass per base pair, a custom DNA fragment length of 106 bp and a stock concentration of 0.256 ng/µl. From these values, we calculated the Cm PCR assay has a detection limit of 3.36 copies.

Real time (qPCR) PCR was carried out using a BioRad CFX machine (Bio-Rad, Hercules, CA). Reactions were run using Qiagen’s QuantiNova Probe PCR Kit (Qiagen, Hilden, Germany) using the kit’s recommended concentrations and cycling conditions. Final Concentration: 1x QuantiNova Master mix, 0.4 µM Primers, and 0.2 µM FAM labeled Probe. Cycling: 95°C 2 min, followed by 40 cycles of 95°C 5 sec, 60°C 30 sec.

All reactions were run in duplicate by loading 5µl of template DNA to 15µl of the reaction mixture. A positive control and negative, no-template control were included in each run. The positive control was a purified PCR amplicon diluted to produce consistent values of Ct 28. A 16s universal bacterial PCR assay using primers 27_Forward and 1492_Reverse was run on all samples to check for DNA extraction and inhibitors. Samples were considered positive if both replicates had similar values and produced a Ct value of less than 40. A sample was called negative if no amplification from the Cm qPCR assay was detected and a positive amplification from the 16s assay was detected.

### Gastrointestinal microbiota analysis

Fecal samples (2 pellets from each mouse) were placed in separate barcoded sample collection tubes containing DNA stabilization buffer to ensure reproducibility, stability, and traceability, and were shipped for DNA extraction, library preparation, and sequencing (Transnetyx Microbiome, Transnetyx, Cordova, TN). DNA extraction was performed using the Qiagen DNeasy 96 PowerSoil Pro QIAcube HT extraction kit and protocol for reproducible extraction of inhibitor-free, high molecular weight genomic DNA that captures the true microbial diversity of stool samples. After DNA extraction and quality control, genomic DNA was converted into sequencing libraries using the KAPA HyperPlus library preparation protocol optimized for minimal bias. Unique dual indexed adapters were used to ensure that reads and/or organisms were not mis-assigned. Thereafter, the libraries were sequenced using the NovaSeq instrument (Illumina, San Diego, CA) and protocol via the shotgun sequencing method (a depth of 2 million 2 × 150 bp read pairs), which enables species and strain level taxonomic resolution. Raw data (in the form of FASTQ files) were analyzed using the One Codex analysis software and database. The One Codex Database consists of approximately 148K complete microbial genomes, including 71K distinct bacterial genomes, 72K viral genomes, and thousands of archaeal, eukaryotic genomes, and mouse gut Metagenome Assembled genomes (MAGs).^46^ Human and mouse genomes are included to screen out host reads. The database was assembled from both public and private sources, with a combination of automated and manual curation steps to remove low quality or mislabeled records. Every individual sequence (NGS read or contig) is compared against the One Codex database by exact alignment using k-mers where k = 31. Based on the relative frequency of unique k-mers in the sample, sequencing artifacts are filtered out of the sample to eliminate false positive results caused by contamination or sequencing artifacts. The relative abundance of each microbial species is estimated based on the depth and coverage of sequencing across every available reference genome.

### Statistical analysis

A Mann-Whitney U (Wilcoxon Rank Sum) test was utilized to determine whether there were statistical differences in survival rates between untreated and treated groups. Survival rates were calculated using the Kaplan-Meier method with statistical differences in survival curves analyzed using the log rank test.^58^ For all statistical analyses performed, a p value equal to or less than 0.05 was considered significant. Data analyses were performed using JMP Pro 17 (SAS Institute Inc., Cary, NC) and survival time is displayed via a Kaplan-Meier curve (SAS 9.4, SAS Institute, Cary, NC).

Time- and antibiotic-dependent effects on the bacterial composition of the gastrointestinal tract were as follows. Shannon and Simpson indices were used to assess α-diversity.^20^ Differences between groups were compared using a one-way repeated measures ANOVA when assessing normally distributed data with equal variance. For data that did not have a normal distribution or data with unequal variance, i.e., the *Amoxi Diet* group, a one-way ANOVA on ranks (Kruskal-Wallis) test was used. Post hoc comparisons were conducted using Tukey’s multiple comparisons test for the former and Dunn’s multiple comparisons test for the latter. When comparing changes over time within the same group, a paired t-test was used for normally distributed data, while for data with a non-normal distribution, i.e., the *Amoxi Diet* group, Wilcoxon matched-pairs signed rank tests were used. All analyses were implemented using GraphPad Prism 9.1.0. For all statistical analyses performed, a *p* value equal to or less than 0.05 was considered significant.

Principal coordinate analysis (PCoA) was conducted on the GM to assess the influence of various treatments on community structure, using both unweighted (Manhattan) and weighted (Bray–Curtis) similarities. To evaluate significance and determine the sources of differences in β-diversity (community membership versus distribution of shared taxa), a one-way permutational multivariate analysis of variance (PerMANOVA) was performed using SciKit-Bio via One Codex software. (One Codex 2024) Where applicable, PerMANOVA p values were controlled for false discovery rates with a two-stage Benjamini, Krieger and Yekutieli procedure using Prism 9.1.0 (Graph Pad, La Jolla, CA)

## Results

### Pilot study

Table 2 details the PCR results for all groups of SW mice. All groups were confirmed Cm PCR positive before treatment. The untreated control group remained PCR positive for Cm at all timepoints (7-, 14- and 58-DPT). Mice treated with *Enro Water* were positive on 7 DPT and not tested further. Mice treated with *TMS Diet-Short* were PCR negative 7 DPT; however, the pooled sample collected from mice in 1 of 2 cages tested positive 14 DPT and thus was considered a treatment failure and not tested further. Mice from all remaining groups were PCR negative at 7-, 14- and 58-DPT.

**Table 2.** *Pilot Study.* Cm PCR results before and after treatment.

| Treatment Group | Pre-Treatment | Days Post Treatment |  |  |
| --- | --- | --- | --- | --- |
|  |  | 7 | 14 | 58 |
| *Control* | + | + | + | + |
| *Amoxi Diet* | + | - | - | - |
| *Doxy Diet* | + | - | - | - |
| *Doxy Water* | + | - | - | - |
| *Enro Water* | + | + | NP | NP |
| *TMS Diet* | + | - | - | - |
| *TMS Diet-S* | + | - | + | NP |
| *TMS Water* | + | - | - | - |

+, positive; -, negative; NP, not performed.

### Cm-infected NSG mice treatment study

#### Survival

All treated NSG mice except those in the *TMS Diet-Long* group were euthanized because of clinical disease at varying times following the end of treatment. Figure 1 displays the survival curves and Table 3 summarizes survival statistics for all treatment groups except *TMS Diet-Long*. Untreated controls began to reach endpoint criteria as early as 14 DPT with all mice in this group euthanized by 33 DPT. Delays to reaching endpoint were treatment specific. NSG mice treated with *Amoxi Diet, Doxy Diet and TMS Water* had significantly prolonged survival compared to the control group. *Amoxi Diet and Doxy Diet* treated mice had the longest mean survival times and with both cohorts’ mice reaching endpoint criteria between 34 and 40 DPT. Mice treated with *TMS Water* reached endpoint criteria between 21 and 34 DPT. *TMS Diet* treated mice did not have significantly prolonged survival compared to the untreated control group. Sick mice exhibited weight loss ≥20% from baseline with hunched postures, dehydration, and lethargy at the time of euthanasia. Those clinical signs were minimal to mild prior to reaching the weight loss benchmark. Mice in the *TMS Diet-Long* group were Cm PCR positive and euthanized 14 DPT as they met positive PCR endpoint criteria. All mice developed a moderate to severe subacute histiocytic and neutrophilic bronchointerstitial pneumonia and/or broncholitis with intralesional chlamydial inclusion bodies (IB) in bronchiolar and alveolar epithelial cells. Cm elementary bodies were also noted in the alveolar spaces. Affected bronchioles exhibited areas of bronchiolar epithelial cell degeneration with single cell necrosis and variable amounts of luminal karyorrhectic debris and degenerate neutrophils, and proteinaceous content.

**Figure 1.**
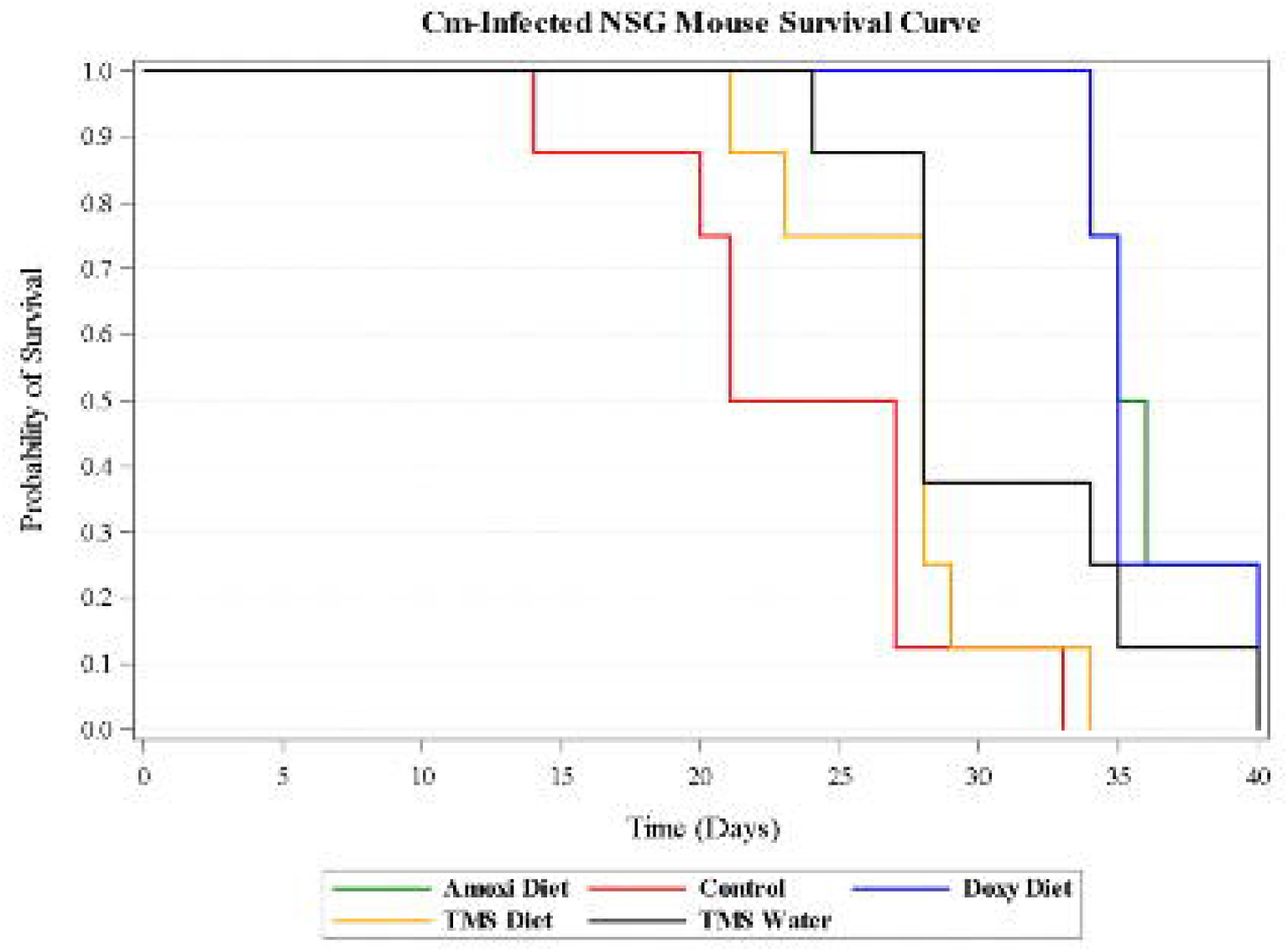
Kaplan-Meier survival curve for untreated and 7-day treatment groups. Day 0 denotes the final day of antibiotic administration.

**Table 3.** Survival data for untreated and 7-day treatment groups in the *Cm-infected NSG treatment study*.

| Treatment Group | Survival Days Post treatment |  |  |  |
| --- | --- | --- | --- | --- |
|  | Mean | SD | Min | Max |
| *Control* | 23.8 | 3.9 | 14 | 33 |
| *Amoxi Diet* | 36.25* | 2.4 | 34 | 40 |
| *Doxy Diet* | 36* | 2.5 | 34 | 40 |
| *TMS Diet* | 27.4 | 3.9 | 24 | 40 |
| *TMS Water* | 30.6* | 5.2 | 21 | 34 |

\* Mean time to endpoint criteria is significantly different ( $P < 0.01$ ) compared to *Control* group.

#### Gastrointestinal microbiota analysis

There were no significant effects of treatment group randomization on diversity based on Shannon (F=0.5626; P=0.6935) and Simpson (F=1.254; P=0.3310) diversity metrics before cohousing or 14 days after acclimatization. Similarly, there were no statistically significant differences in β-diversity metrics when assessing fecal bacterial community diversity or its’ composition (Manhattan [F=1.512; p=0.087]; Bray-Curtis [F=1.512; p = 0.086]; Figure 2). Cohousing with a C mouse significantly altered the NSG cage mates’ microbiota. Comparison of richness and evenness between all NSG mice before (0 DPC) and after (14 DPC) cohousing revealed significant changes in both Shannon and Simpson diversity indices (P=0.0005; P=0.0003, respectively; Figure 3 A, B). PCoA plots revealed a clear separation in fecal bacterial communities before and after cohousing, and PerMANOVA tests revealed statistical differences between the microbial communities present at 0 and 14 DPC (Manhattan [F=6.293; p=0.001]; Bray-Curtis [F= 6.293; p = 0.001]; Figure 3 C). Comparison of richness and evenness before (14 DPC) and after (7 DPT) *Doxy Diet* treatment revealed a significant effect on both Shannon (P=0.0458) and Simpson (P=0.0401) diversity. One-way PerMANOVA testing confirmed these effects using both weighted and non-weighted similarity metrics. *TMS Water* also had a significant effect on the microbiota with significance in Shannon (P=0.0171), but not Simpson diversity. Significant differences in Shannon and Simpson diversity before (14 DPC) and after (7 DPT) antibiotic treatment were not observed in the TMS Diet, Amoxi Diet, or Control groups. Manhattan distance and Bray-Curtis dissimilarity (Table 5, Figure 4) of the fecal bacterial communities of all groups were significantly different when comparing them at 14 DPC and 7 DPT.

**Figure 2.**
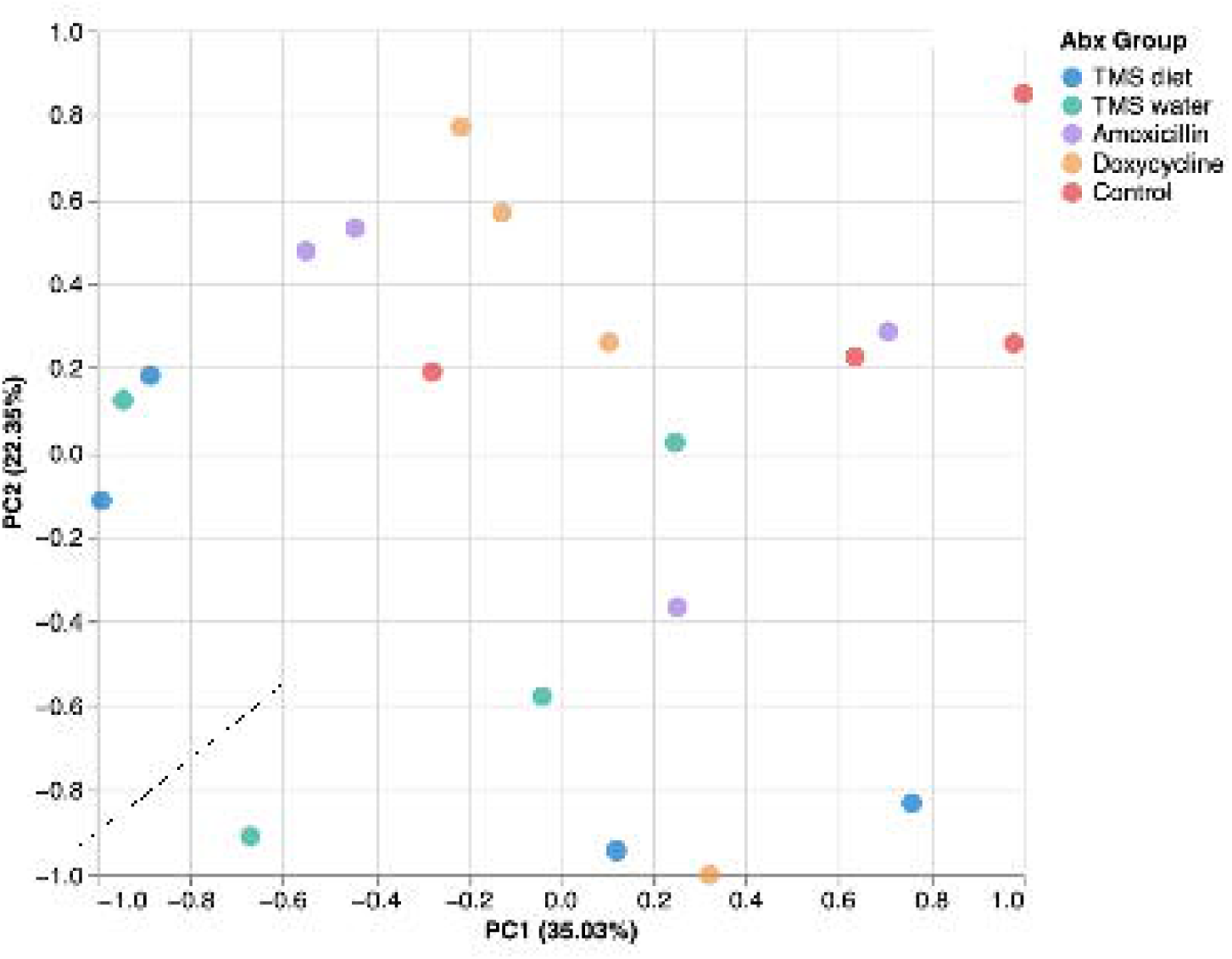
PCoA of samples collected from NSG mice (n = 20) after randomization, but before cohousing with C mice using Bray-Curtis Similarity (A). Colored datapoints denote untreated mice destined for a specific treatment. Manhattan Distance was not included due to similarity of the plot to Bray-Curtis Similarity

**Figure 3.**
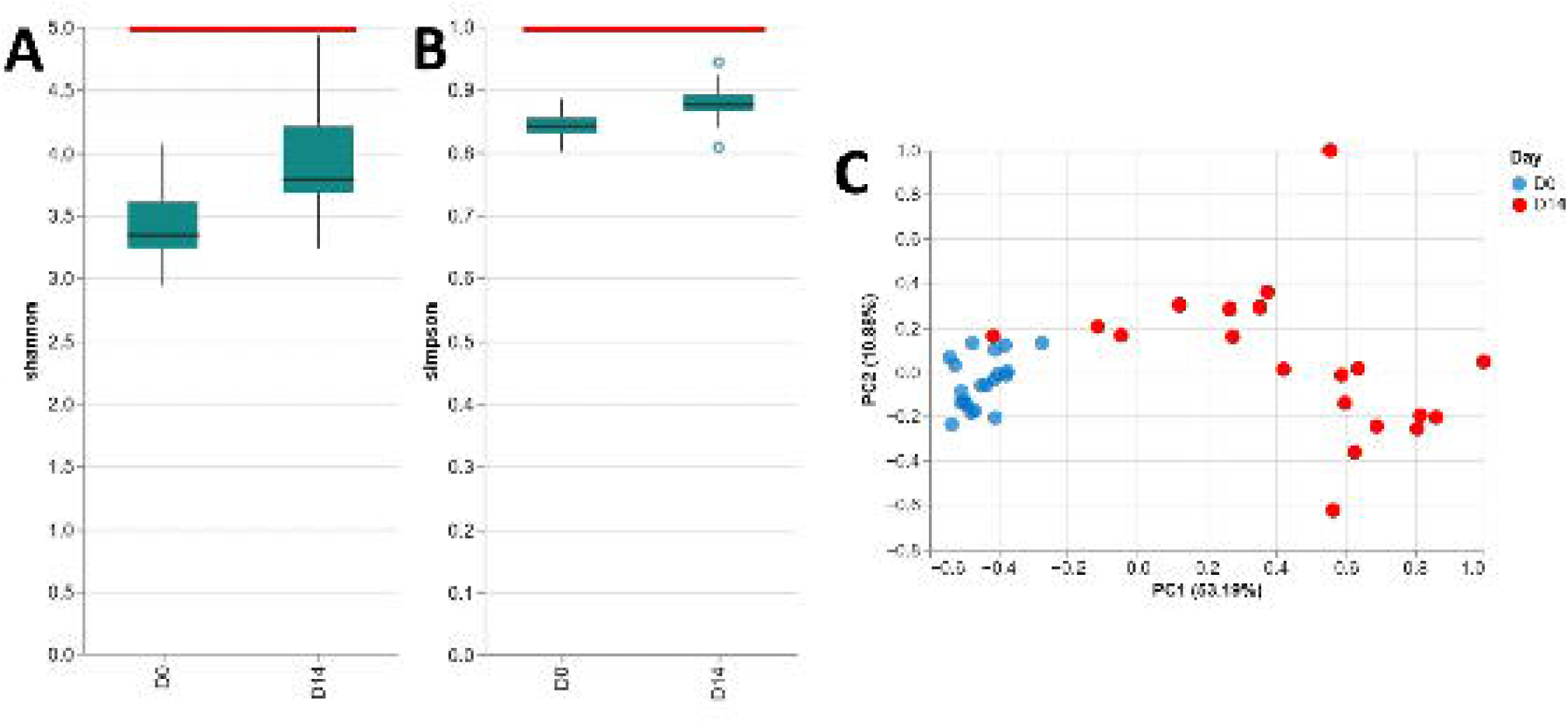
α-diversity and PCoA of samples collected from NSG mice (n = 20) after randomization and cohousing, but before antibiotic treatment using Shannon Index (A), Simpson Index (B), and Bray-Curtis Similarity (C). Manhattan Distance was not included due to similarity of the plot to Bray-Curtis Similarity. Lines Indicate significant (p < 0.05) time-dependent differences as determined via paired t test.

**Figure 4.**
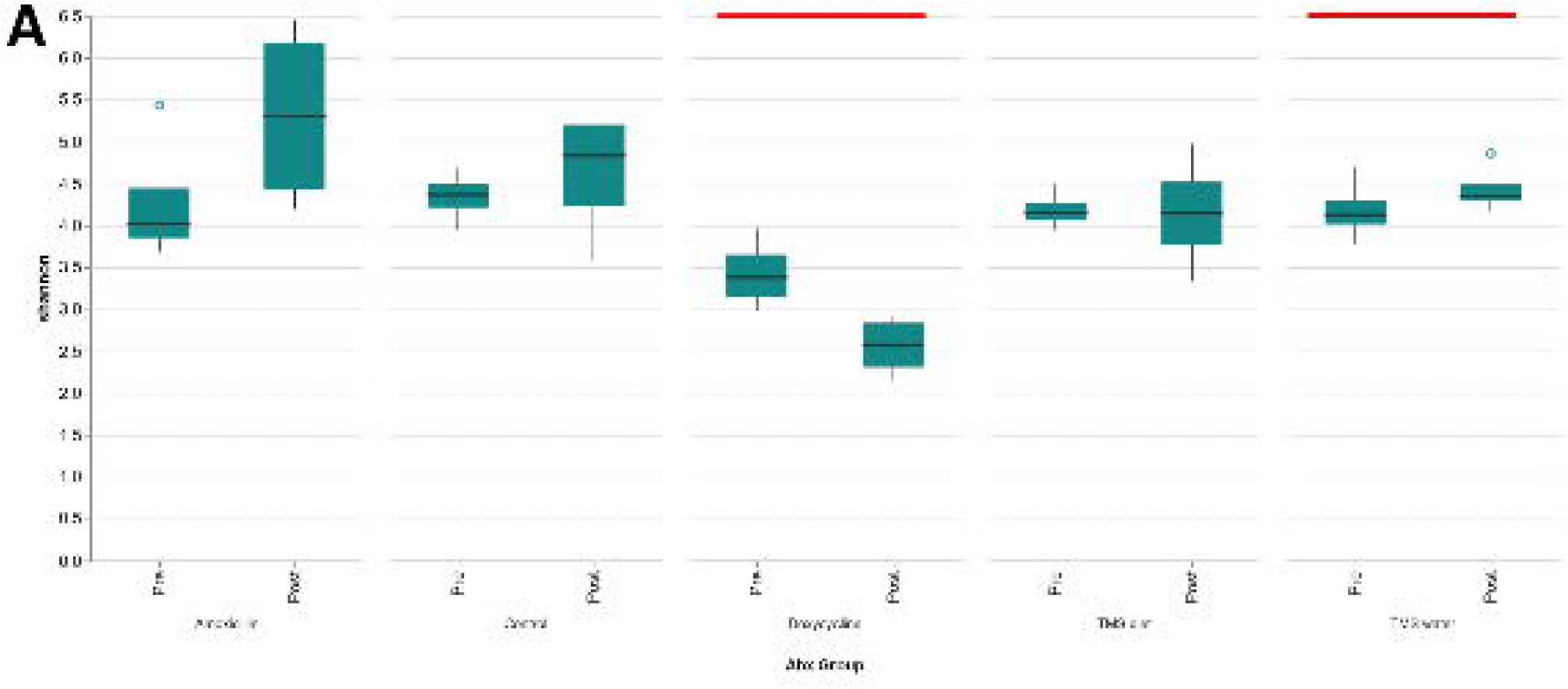
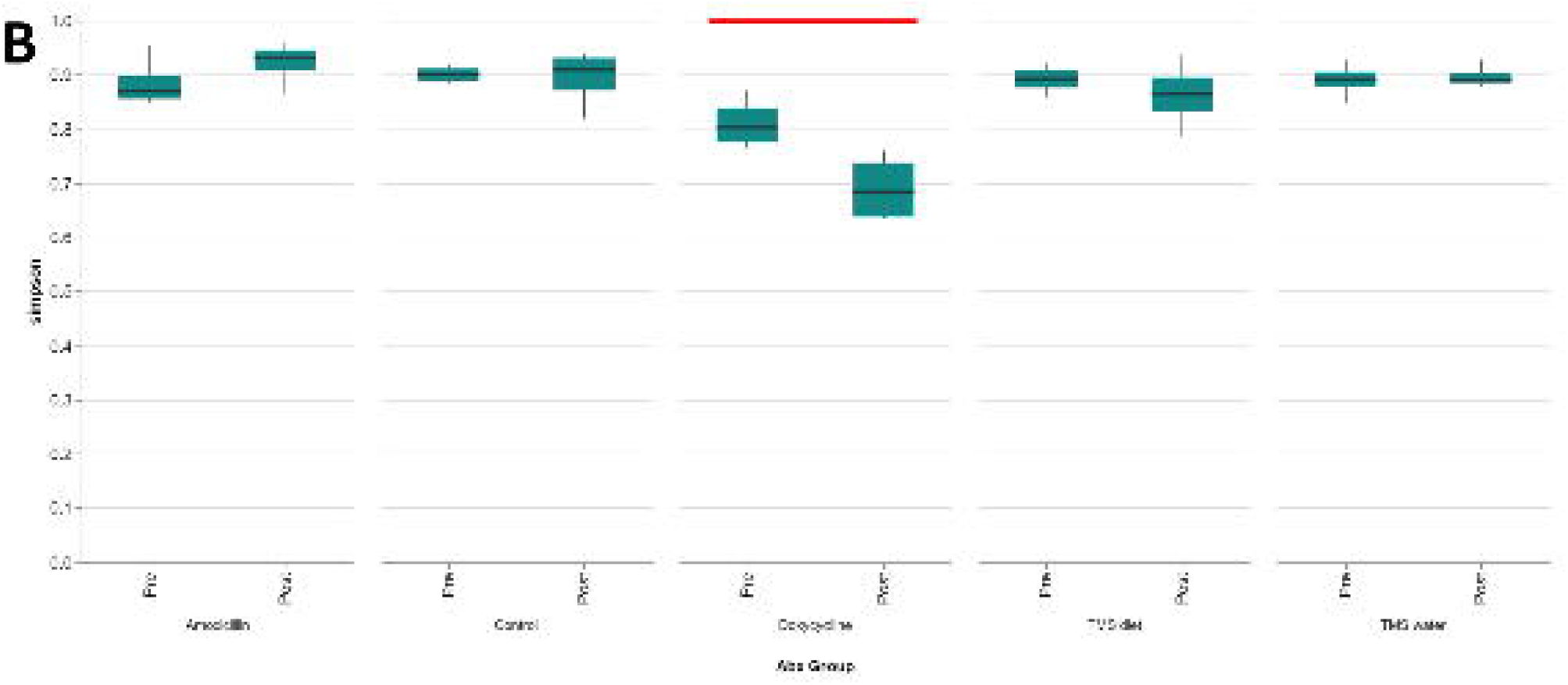
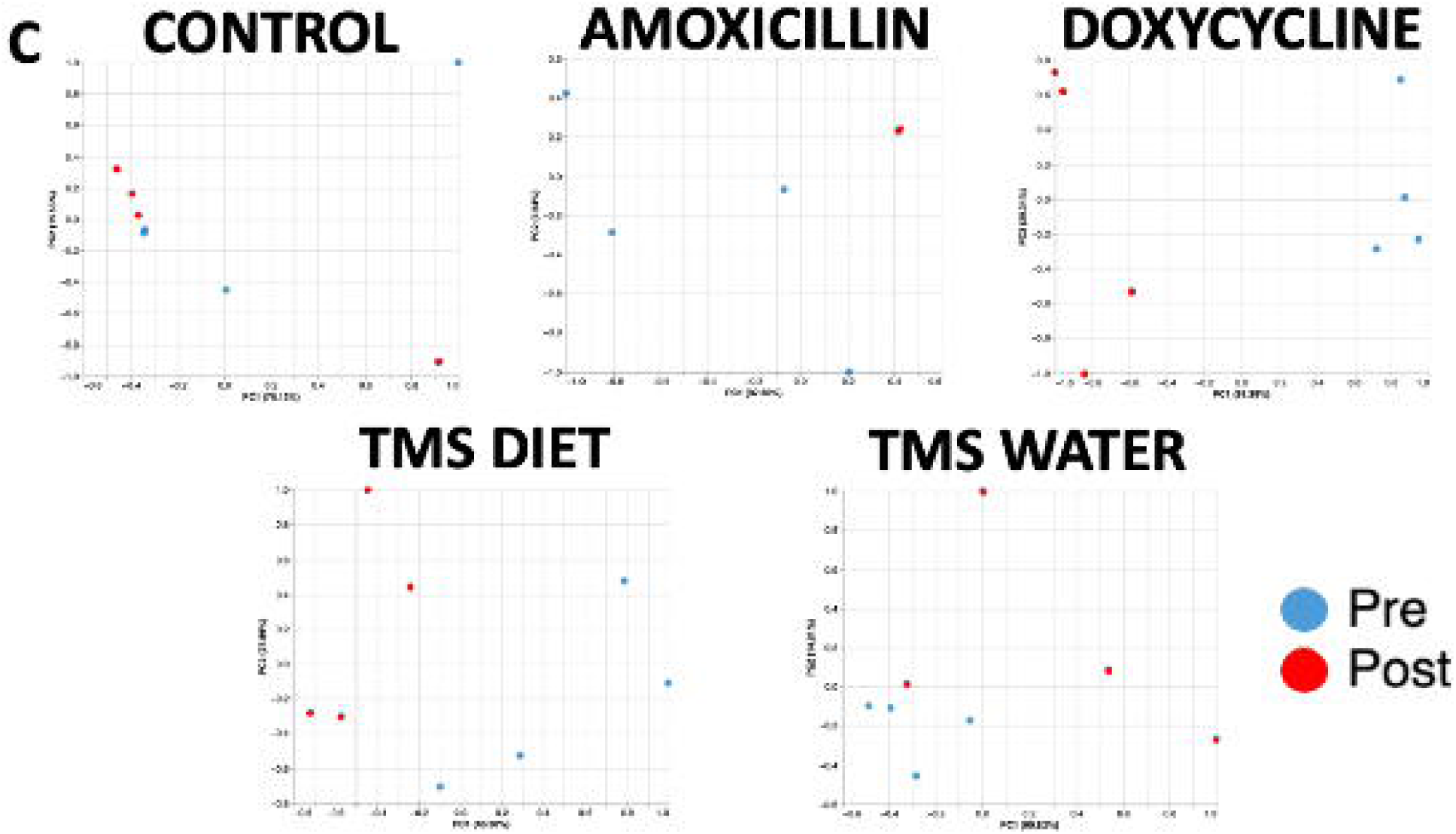
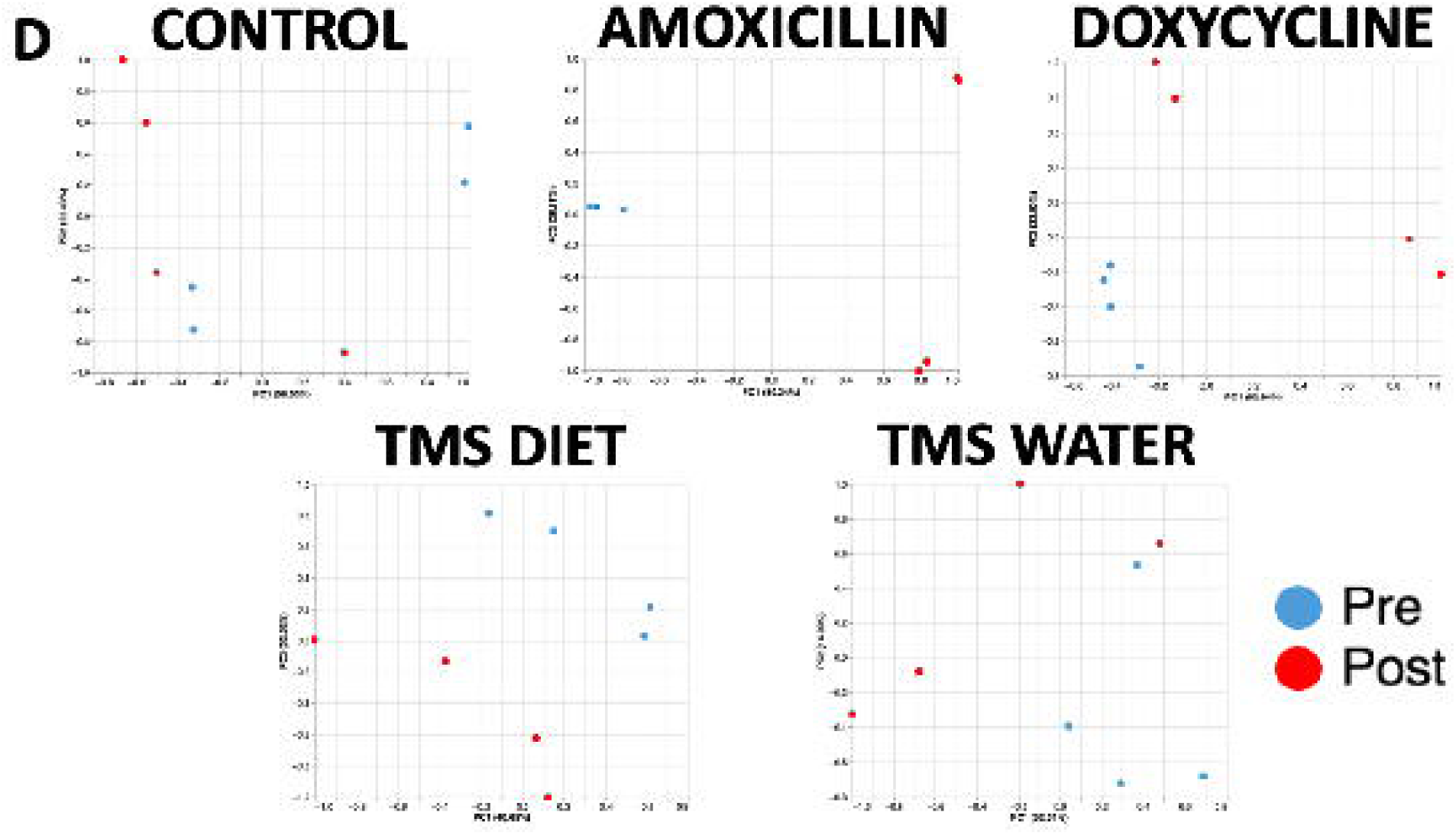
α-diversity and PCoA plots for samples collected from each group (n = 4) before (pre) and on the last day of (post) antibiotic treatment using Shannon Index (A), Simpson Index (B), Manhattan Distance (C) and Bray-Curtis Similarity (D). Red lines indicate significant (p < 0.05) time- and antibiotic-dependent differences as determined via paired t test or Wilcoxon matched-pairs signed rank tests.

### Inbred immunocompetent & NOD.SCID mouse study

All groups were confirmed Cm PCR positive before treatment. The untreated control group remained PCR positive, and all treated groups were Cm PCR negative for up to 40 DPT (Table 4). Antibiotic treated mice did not display clinical signs throughout the study duration. Similarly, untreated NOD.SCID mice did not develop clinical signs despite remaining PCR positive. However, 1 of 2 untreated NOD.SCID mice had a marked bronchointerstitial pneumonia affecting the right pulmonary lobes and a mild suppurative rhinitis at necropsy. Chlamydial IBs were noted in affected bronchiolar epithelial cells; these cells also demonstrated MOMP immunolabeling. The lungs from the second untreated NOD.SCID mouse had minimal alveolar histiocytosis and peribronchiolar lymphocytic infiltrate; no IBs nor MOMP immunolabeling was detected. Chlamydial IBs and MOMP immunolabeling were occasionally detected in the colonic epithelium of both untreated NOD.SCID mice. There were no IBs nor MOMP immunolabeling detected in any of the tissues examined in the treated C, B6, and NOD.SCID mice.

**Table 4.** Cm fecal PCR results in the *Inbred immunocompetent and NOD.SCID mouse study*.

| Treatment Group | Pre-Treatment | Days Post Treatment |  |  |
| --- | --- | --- | --- | --- |
|  |  | 7 | 14 | 40 |
| **Control** | + | + | + | + |
| **C** | + | - | - | - |
| **B6** | + | - | - | - |
| **NOD SCID** | + | - | - | - |

+, positive; -, negative.

## Discussion

*Chlamydiaecae* have a unique life cycle, a wide host range, diverse tissue tropism, and treatment resistance in humans, mice, goats, sheep, cattle, cats, koalas, swine, guinea pigs and over 130 avian species.^9,22,31,45,63^ All chlamydial species exhibit a biphasic life cycle, composed of a metabolically inert infectious elementary body (EB) and a metabolically active reticulate body (RB).^18,22,52^ When the EB is internalized by a phagosome in the host mucosal cell, an IB is created.^18,52^ Within the IB, the EB differentiates into a RB and begins to replicate until cellular lysis or extrusion of newly generated infectious EBs occurs. Released EBs infect nearby cells or are excreted and infect naive hosts.^18,52,63^ A deviation from this normal life cycle can occur when RBs are exposed to environmental, immunologic, or pharmaceutical stressors.^9,13,16,41,51,56,73^ Under stress, aberrant bodies (ABs) may be formed and enter a persistent antibiotic-resistant state. ABs have an increased resistance to stress allowing the bacterium to remain viable although ABs are not infectious.^7,9,13,41,51,56,72,73^ Interestingly, upon removal of the stress, ABs undergo redifferentiation, a mechanism that remains poorly understood, transitioning back into RBs and subsequently into infectious EBs, thereby completing the life cycle and perpetuating transmission.^50,51,52^ This makes chlamydia challenging to eradicate without a highly efficacious treatment regimen.

Since initiating testing for Cm in mice in 2021, we have successfully treated small numbers of many unique strains of mice, often with unknown and likely varying immunocompetencies, imported from other research colonies with TMS provided in the drinking water. However, a controlled study to identify safe, easy to administer, effective therapeutics for Cm that would avoid undesirable effects like chlamydial persistence or that would cause abrupt changes in microbiota was necessary before considering treating colonies of Cm-infected mice. Further, as many genetically engineered and mutant mouse (GEM) strains are immunodeficient, we felt compelled to evaluate treatment effectiveness in mice with a significant immunodeficiency to understand the likelihood of treatment failure in mice that are immunocompromised.

A pilot study was conducted initially using various antibiotics, known to be effective against chlamydial species, and delivery methods by treating retired, Cm-infected SW sentinels. All antibiotic formulations used in the pilot study were selected based on clinical and experimental experience. Amoxicillin, TMS and doxycycline medicated feed are commercially available and routinely used in our vivaria. We included doxycycline medicated water because many laboratories use it for research purposes but avoided amoxicillin in water because we never use it in our vivaria since compounded feed is commercially available and considerably easier to administer. We also suspected amoxicillin would not be a preferred antibiotic since it was expected to significantly impact the microbiota. Following treatment, we cohoused highly immunodeficient NSG mice, which are detrimentally affected by Cm infection, with the treated cage mates to increase the sensitivity of detecting treatment failures.^66^ Most of the antibiotics evaluated were successful when administered for 7 days except for enrofloxacin or a short (3 day) course of TMS. Enrofloxacin, a bactericidal fluoroquinolone failed despite its effectiveness against both gram-positive and gram-negative bacteria, its ability to achieve high intracellular concentrations and chlamydial treatment successes in other species.^27,53^ Feces from mice housed in 1 of 2 cages treated with TMS for 3 days tested positive for Cm 14 DPC even though the mice had been tested negative by PCR on 7 DPT. Failure was likely a result of the short duration of treatment allowing residual EBs, RBs and/or ABs to remain viable. Although TMS is bactericidal, the formation of ABs offering protection from the antibiotic has been reported in previous studies in which TMS was administered.^26,30^ TMS has been shown to have a short half-life in mice, so this in conjunction with the short treatment duration may have allowed animals to harbor extremely low levels of viable Cm post-treatment.^4,25^ These levels of Cm were likely below the detection threshold of the PCR assay. Subsequent exponential growth over the following 7 days led to the proliferation of Cm and detection of the bacterium by PCR 14 DPT. Another possibility for treatment failure was the potential for autoinfection via coprophagy with the ingestion of feces containing viable Cm EBs led to reinfection. This scenario is least likely given that a complete cage change was performed for all groups at the time of antibiotic removal. Although the reason for treatment failure could not be definitively determined, these treatment regimens were not investigated further.

We then treated Cm-infected NSG mice for 7 days with 1 of the 4 antibiotics which were used successfully in the pilot study; *Doxy Water* was not pursued because doxycycline compounded feed is available commercially and would be considerably easier to administer to large numbers of mice. Unfortunately, none of the treatment protocols eliminated Cm. The survival data indicated that the antibiotic protocols had some protective effects, prolonging survival in mice which otherwise would have succumbed earlier to Cm for varying periods. We then decided to evaluate a 21-day treatment protocol in NSG mice. We only used TMS compounded feed because it was effective in the pilot study, is commercially available from multiple manufacturers, provides a more stable ingested dose over time, and is easier to administer to a large colony in contrast to medicated water.^42^ Further, administering TMS in the drinking water would be challenging in vivaria that make extensive use of automated watering systems. Ultimately, the 21-day treatment protocol trial failed. Thus, our results demonstrated that while select antibiotics can delay clinical signs and prolong survival, at least some immune pathways lacking in the NSG model are necessary to eradicate following antibiotic therapy.

Failure to treat Cm-infected NSG mice prompted us to consider treatment efficacy in mice which may not be as immunocompromised as the highly immunodeficient NSG strain and would better model Cm-infected genetically engineered strains which may have an altered immune responsiveness, that may be resident within our colonies. In addition, we chose to evaluate treatment efficacy in B6 and C mice because B6 mice have a Th-1 biased immune response while C mice exhibit a Th-2 biased immune response and these are the most commonly used inbred strains and serve as the genetic background for many GEM strains.^69^ NOD.SCID mice were chosen for treatment evaluation because they are significantly immunocompromised, although their innate immune system deficiencies are not as profound. Because of their moderate immunodeficiency we elected to treat NOD.SCID mice for 21 days while the immunocompetent B6 and C mice were treated for 7 days. Cm was successfully eradicated from NOD.SCID mice as they remained PCR negative up to 40 DPT, all tissues evaluated at necropsy were histologically normal, MOMP immunolabelling was not observed, and NSG contact sentinels remained clinically healthy and PCR negative up 40 DPT. The question remains as to whether a reduced treatment duration, perhaps as short as the 7-day duration used successfully for the immunocompetent B6 and C strains, would achieve a similar result. This remains an important question if considering treating large colonies or entire facilities as there are likely resident strains that have unrecognized immunodeficiencies. Importantly and relevant to the risk of inadequately treated strains remaining a risk of reinfection, we recently reported that neither contaminated caging nor barrier-level husbandry methods led to intercage transmission of Cm.^47^

Interestingly, although both untreated control NOD.SCID mice were PCR positive at 40 DPT, only 1 of 2 developed a subclinical Cm-associated pneumonia, suggesting that the presence of IL-2rγ-dependent immune pathways lacking in NSG mice might be essential for the control of Cm infection. We hypothesize that the preservation of NK cells and other innate lymphoid cells in NOD.SCID mice allowed for the phagocytosis of viable Cm and the production of the various cytokines necessary for successful Cm clearance.

A secondary aim of the *NSG Treatment Study* was to determine if changes in the GM that are associated with antibiotic use can be minimized with the selection of a specific antibiotic. Not surprisingly, changes in the GM occurred in NSG mice when exposed to C mice. While both NSG and C mice used in this study originated from the same vendor, strain specific and age dependent microbiota diversity exists even in mice housed in the same room.^39,48^ GM analysis confirmed different antibiotics and perhaps different formulations of a single antibiotic (i.e., *TMS Diet* and *TMS Water*) can have differing effects. Our untreated control mice showed the least changes in GM, as determined by *p* and F value significance (Table 5). While there were no significant changes in α-diversity after cohousing with infected C mice, significant changes in β-diversity were detected in these mice, which was to be expected as microbial communities are constantly in flux.^2,35^

**Table 5.** Results of one-way PerMANOVA controlling for false discovery rates in p values with a two-stage Benjamini, Krieger and Yekutieli procedure to detect differences in fecal community structure between pre-treatment (Pre) and and on the last day of treatment (Post) samples from NSG mice receiving antibiotics. (n = 4)

| Treatment | Similarity | Pre vs Post |  |
| --- | --- | --- | --- |
|  |  | *P* | **F** |
| *Control* | Manhattan | 0.044 | 1.43 |
|  | Bray-Curtis | 0.046 | 1.43 |
| *Amoxi Diet* | Manhattan | 0.008 | 8.91 |
|  | Bray-Curtis | 0.009 | 5.66 |
| *Doxy Diet* | Manhattan | 0.008 | 8.356 |
|  | Bray-Curtis | 0.009 | 8.356 |
| *TMS Diet* | Manhattan | 0.009 | 7.457 |
|  | Bray-Curtis | 0.009 | 7.458 |
| *TMS Water* | Manhattan | 0.008 | 4.134 |
|  | Bray-Curtis | 0.009 | 4.134 |

Oral administration of TMS in the drinking water for 7 days resulted in significant time-dependent alterations in microbiota richness and β-diversity when comparing the GM before and after treatment. This is inconsistent with the previous report by Korte et al., who showed that oral administration of TMS in the drinking water for 14 days caused negligible effects on the GM of mice on a B6 background.^33^ Both the use of NSG mice, which are known to exhibit more unstable microbiota compared to wild type mice, and the longer treatment protocol may have contributed to the different result.^2,33^ Conversely, mice treated with *TMS Diet* did not exhibit significant changes in α-diversity, but did have significant β-diversity changes. The most likely explanation is that although the overall diversity within each sample remained consistent, the selective eradication of bacterial species by the antibiotic allowed the GM to become reconstituted differently in each mouse by species that colonized the vacant ecological niches. Interestingly, *TMS Diet* treated mice did not exhibit significant effects on α-diversity despite providing a higher estimated daily ingested dose than *TMS Water*. It is unclear why an antibiotic at a lower dose would have a more significant effect, but TMS-compounded diets have less dosing variability than TMS in water.^42^ Trimethoprim has low water solubility (0.4 mg/ml) while sulfamethoxazole is “practically insoluble”, leading to particulate aggregation and sedimentation in the bottle.^42^ As the water bottle sits at an angle with the sipper tube pointed down, the sedimentation settles in close proximity to the sipper tube and would likely have resulted in the mice ingesting a considerably larger dose than estimated. In contrast to the other antibiotics evaluated in this study, TMS’ lessor impact on the gastrointestinal microbiota may be because it is absorbed in the upper gastrointestinal tract and is primarily eliminated via renal excretion with minimal hepatic metabolism.^33^

*Doxy Diet* had the most significant effects on both α and β diversities. This was not surprising as doxycycline is recognized to induce dysbiosis, has immunomodulatory effects, and may even contribute to developmental defects in offspring.^10,28,49^ This effect, as well as the relatively large number of GEM strains with Tet-on/Tet-Off doxycycline inducible genes, makes *Doxy Diet* an undesirable choice for treating large numbers of mice. *Amoxi Diet* did not induce significant changes in α diversity in this study, which contrasts with other studies that have shown significant microbiota alterations in mice treated with amoxicillin at doses both higher and lower than what we administered. ^38,65^ Overall, based on its limited effect on the microbiota and its’ effectiveness eradicating Cm, TMS would be the preferred antibiotic for treating Cm positive mice. While our results suggest doxycycline and amoxicillin are equally effective, they have been reported to cause significant changes to the GM.^32,37,38,65^ Importantly, our analysis was limited by sample size and the fact that we only treated Cm infected female mice thus cannot assess the effect that gender may have on eradication or microbiota results. However, the similarity in the microbiota changes resulting from administration of TMS in both feed and water as well as with published reports, provides compelling evidence that TMS is likely to have less impact on the microbiota than the other antibiotics evaluated.

In conclusion, we demonstrated that TMS can eradicate Cm from wild type immunocompetent strains and stocks following 7 days of treatment, and from a moderately immunocompromised following 21 days of therapy. *TMS Diet* would be our preferred treatment as it would have the least impact on the microbiota and is easy to obtain and administer. Treatment efficacy should be evaluated by confirmatory PCR testing.

## Abbreviations

Cm: *Chlamydia muridarum*
GEM: genetically engineered mouse
NSG: NOD.Cg-*Prkdc^scid^ Il2rg^tm1Wjl^*/SzJ
SW: Tac:SW
C: BALB/cJ
B6: C57BL/6J
GM: gastrointestinal microbiota
TMS: trimethoprim and sulfamethoxazole
NOD.SCID: NOD.Cg-*Prkdc^scid^Il2rg^tm1Wjl^*/SzJ
IHC: Immunohistochemistry
MOMP: major outer membrane protein
EB: elementary body
RB: reticulate body
AB: aberrant body
IB: inclusion body
DPT: days post-treatment

## Acknowledgements

The authors would like to thank Simona Bekker and John D’Allara for their technical assistance during sample handling and shipment, Fabricio Munoz and Marcia Lewis for their assistance with breeding of the NSG mice used in this study. We also thank the Laboratory of Comparative Pathology staff for their technical assistance with histology and IHC.

